# Optical single-channel recording of CRAC channels with HaloTag and a Ca^2+^-sensitive ligand

**DOI:** 10.64898/2026.05.08.723778

**Authors:** Harsharan Dhillon, Richard S. Lewis

**Affiliations:** Neurosciences Program and Department of Molecular and Cellular Physiology, Stanford University School of Medicine, Stanford, CA 94305

## Abstract

Following ER Ca^2+^ depletion, Ca^2+^ release-activated Ca^2+^ (CRAC) channels are activated by STIM1 at ER-plasma membrane junctions. The restricted localization and low conductance of the CRAC channel (<40 fS) precludes single-channel recordings, limiting studies of CRAC channel gating. Here we describe an optical approach to characterize the gating of HaloTag-fused Orai1 channels labeled with JF646-BAPTA, a Ca^2+^-sensitive fluorescent dye. While Ca^2+^ influx through single channels generates fluorescence fluctuations, identifying true gating events is complicated by stochastic transitions of JF646-BAPTA to a non-fluorescent state. To overcome this, we combine TIRF microscopy with whole-cell voltage clamp to control the driving force for Ca^2+^ entry. We show the open channel intensity at -100 mV reflects Ca^2+^ saturation of the dyes on each channel, while the closed-channel intensity is defined by the fluorescence at +30 mV, where influx is absent. True gating events can be identified from transitions between the open- and closed-channel levels, distinguishing them from transitions to a non-fluorescent state. We describe the gating behavior of CRAC channels activated by STIM1 after store depletion. Dwell time distributions indicate at least two open and closed states with durations of 0.1 to several seconds, with most channels having an open probability of ≥0.7. We also detect ‘silent’ channels that colocalize with STIM1 but show no activity over tens of seconds, a population that would be undetectable by whole-cell electrophysiology alone. This method offers an approach to explore CRAC channel gating mechanisms and may be applicable to other Ca^2+^- permeable channels not amenable to patch-clamp techniques.

## INTRODUCTION

Store-operated Ca^2+^ entry (SOCE), mediated by Ca^2+^ release-activated Ca^2+^ (CRAC) channels, is a major signaling pathway involved in a wide range of cell functions including gene expression, cell motility, and secretion (Prakriya and Lewis, 2015). Loss-of-function mutations have shown that CRAC channels are critical for activation of the immune response as well as muscle development and function and other essential physiological processes (Emrich et al., 2022; Vaeth et al., 2020). The CRAC channel comprises two components: Orai1, the pore-forming subunit in the plasma membrane (PM), and STIM1, an ER resident protein that senses Ca^2+^ in the lumen of the endoplasmic reticulum (ER).

Depletion of the ER Ca^2+^ store by stimulation of cell surface receptors activates a conformational change in STIM1, causing it to accumulate at ER–PM contact sites where it binds and traps diffusing Orai1 channels in the PM and opens them to trigger Ca^2+^ influx (Prakriya and Lewis, 2015). Ca^2+^ entry through Orai1 acts both globally through elevation of [Ca^2+^]_i_ throughout the cytosol as well as locally at ER-PM junctions. Local signaling has been implicated in modulation of the PM Ca^2+^-ATPase (Bautista and Lewis, 2004), adenylate cyclase (Cooper, 2015), and local uptake by SERCA Ca^2+^ pumps (Courjaret et al., 2025), as well as activation of calcineurin to dephosphorylate the transcription factor NFAT (Kar et al., 2021), enabling its translocation to the nucleus where it regulates a large variety of responsive genes (Feske et al., 2001).

Much of the progress made in understanding the mechanisms underlying CRAC channel activation have relied on whole-cell studies with native cells or heterologous overexpression of STIM1 and Orai1. This approach simplified the tracking of STIM1 and Orai1 movements during activation and characterization of CRAC Ca^2+^ current in whole-cell recording, revealing a distinctive biophysical fingerprint of extremely high Ca^2+^ selectivity and low unitary conductance. However, the stochastic activity of single channels is obscured by ensemble averaging of hundreds to thousands of channels during whole-cell recording. Experimental evidence and modeling suggest that the number of channels at each junction is actually quite low, in the range of 1-5 depending on cell type (Hogan, 2015; Shen et al., 2021). Thus, resolving single-channel behavior is key to understanding local signaling at ER-PM junctions. Single-channel recording is also critical for assessing the open probability and kinetic behavior of the channel, which together with mutagenesis and structural studies can be applied to elucidate mechanisms of gating and channel modulators. Unfortunately, the extremely small conductance of the CRAC channel poses a major obstacle to single-channel recording with patch-clamp techniques. Fluctuation analysis of whole-cell currents has yielded estimates of 20-40 fS, roughly 100-fold smaller than that of voltage-gated Ca^2+^ channels, and too small to be detected against a background of thermal noise (Zweifach and Lewis, 1993; Prakriya and Lewis, 2006).

An alternative approach is to optically monitor Ca^2+^ flux through single channels by attaching a Ca^2+^-sensitive fluorescent probe directly to the channel. Because Ca^2+^ concentration falls steeply with distance as it diffuses from the mouth of the pore, a tethered indicator can in principle be made sensitive only to Ca^2+^ entering through the channel to which it is attached, effectively reporting diffraction-limited single-channel activity across large areas of membrane. In a pioneering study, Dynes et al. recorded optical activity of the Ca^2+^-sensitive fluorescent proteins G-GECO1 or G-GEC01.2 fused to Orai1 (Dynes et al., 2016). Single Orai1 channels in cells coexpressing a low level of the soluble CRAC activation domain (CAD) from STIM1 showed infrequent single flickers lasting several hundred ms and longer transients lasting 1 to several s. Higher levels of CAD as well as STIM1 in store-depleted cells evoked oscillatory responses without the expected binary signals associated with channel opening and closing. While these studies were a significant advance in monitoring single Orai1 channel behavior, it is difficult to extract open probability or open and closed durations from the data, particularly from the oscillatory responses of single channels activated by STIM1. One reason for this may be that the properties of these Ca^2+^ -indicator GECI proteins, including the GECO and GCaMP series, are not ideal for this application: the relatively slow on and off rates of GECIs (10’s to 100’s of ms) and moderate brightness limit temporal resolution, GECIs are prone to photobleaching which limits recording duration, and the blue excitation light for GCaMPs generate cell autofluorescence, reducing the signal to noise ratio (SNR) of the emitted signal.

A new approach based on Ca^2+^ -sensitive fluorogenic dyes bound to HaloTag promises to overcome some of the limitations of GECIs for monitoring single-channel activity. JF646-BAPTA combines the Ca^2+^ chelator BAPTA with the Si-rhodamine dye JF646 (**Fig. S1A**), and on covalent attachment to HaloTag it exhibits Ca^2+^-sensitive fluorescence with a dynamic range of ∼5 and a K_d_ of 150 nM (Deo et al., 2019). Compared to proteins in the GCaMP series, JF646-BAPTA is brighter, less prone to bleaching, less environmentally (pH) sensitive, has faster Ca^2+^ binding and unbinding kinetics, and because of its far-red excitation and emission wavelengths produces less autofluorescence (Deo et al., 2019). In its first application as a reporter of channel activity, JF646-BAPTA was used to monitor the activity of mechanosensitive PIEZO1 channels fused to HaloTag in hiPSC-derived cells and organoids (Bertaccini et al., 2025). The results revealed the spatial distribution of active channels, opening a new window on how mechanical stimuli are transduced in different cell types and during development. Remarkably, rapidly flickering fluorescence signals resembling single-channel openings and closings were observed, suggesting that JF646-BAPTA could be used to study gating at the single-channel level.

Here we introduce a method for monitoring the activity of single Orai1 channels fused with HaloTag and labeled with JF646-BAPTA. We demonstrate that the dye undergoes reversible transitions to a non-fluorescent state (‘blinking’) which can confound identification of single channel gating events. To overcome this problem, we describe a strategy for detecting single Orai channel activity and measuring true open-closed transitions based on the simultaneous response of several dyes attached to the channel. Using this strategy, we quantify the duration of open and closed states of channels in contact with STIM1 and measure their open probability. This method should be generally applicable to a variety of Ca^2+^ -permeable channels in addition to Orai to reveal gating kinetics and the spatial distribution of channel activity underlying a range of physiological processes.

## RESULTS

### Labeling Orai1 with HaloTag and a Ca^2+^-responsive HaloTag ligand

HaloTag was appended via a 14-residue linker to the C-terminus of Orai1, resulting in a hexameric channel with six HaloTags. Methods for labeling the HT with JF646-BAPTA/AM in different cell systems have varied widely, from 1 µM overnight (Deo et al., 2019) to 500 pM for 15 min (Bertaccini et al., 2025). To evaluate labeling strategies, we reacted free Orai1-HaloTag sites with the membrane-permeant dye JF-552 after incubation of cells coexpressing GFP-STIM1 with JF646-BAPTA/AM. Both dyes were imaged by TIRF of the cell footprint after treatment with TG to deplete ER Ca^2+^ and induce STIM1/Orai1 puncta (**Fig. S1B**). Following overnight exposure to 1 µM JF646-BAPTA/AM, JF-552 clearly labeled free sites not occupied by JF646-BAPTA (**Fig. S1B**, left column). In contrast, 1 µM JF646-BAPTA/AM in 0.02% pluronic F-127 achieved complete labeling of all sites in 1 h, as shown by the lack of subsequent labeling with JF-552 (**Fig. S1B**, right column). Cell background fluorescence in the JF646-BAPTA channel was not detectably increased by the labeling protocol, presumably because JF646 is highly fluorogenic, increasing 21-fold in fluorescence upon binding to the HaloTag (Grimm et al., 2015).

Although fusing GFP to the C-terminus of Orai1 does not significantly perturb its biophysical properties (Mullins et al., 2016; Yen and Lewis, 2018), HaloTag is somewhat larger, and because STIM1 also binds to the C-terminus of Orai1 it was important to test whether it is similarly benign. When coexpressed with GFP-STIM1, Orai1-HaloTag colocalized with STIM1 after ER Ca^2+^ depletion with TG (**Fig. 1A**). Colocalization was precise, as shown by line scans through several puncta (**Fig. 1B**). In cells expressing GFP-STIM1 and Orai-HaloTag/JF646-BAPTA the time course of I_CRAC_ activation by EGTA in the whole-cell pipette solution also appeared normal (**Fig. 1C**). The current-voltage relation of I_CRAC_ showed characteristic inward rectification and a lack of outward current above +50 mV, indicating that the pore structure is not significantly altered by the attachment of HaloTag (**Fig. 1D**). Together, these results confirm that the Orai1-HaloTag chimera labeled with JF646-BAPTA maintains characteristic biophysical and physiological properties of the CRAC channel.

**Figure 1.**
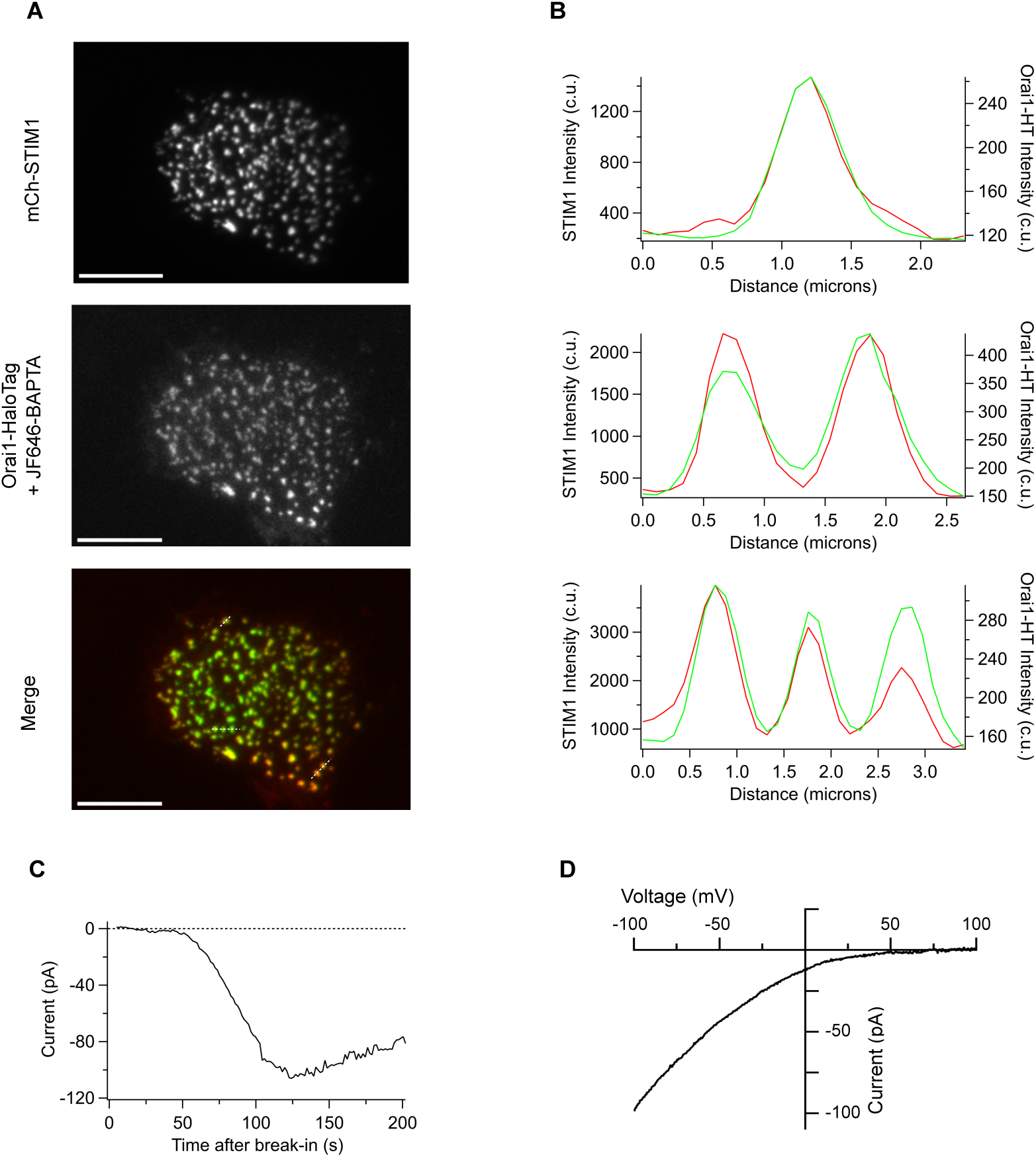
HaloTag labeling of Orai1 does not perturb Orai1 function. **(A)** TIRF images of TG-treated (store-depleted) HEK293 cells expressing Orai1-HaloTag and mCh-STIM1 after loading with JF646-BAPTA. The merged image shows colocalization of mCh-STIM1 (green) and Orai1-HaloTag (red). **(B)** Line scans of mCh-STIM1 (green) and Orai1-HaloTag (red) from selected puncta in the merged image from A. **(C)** Induction of I_CRAC_ in a HEK293 cell expressing mCh-STIM1 and Orai1-HaloTag after break-in to the whole-cell recording configuration with EGTA in the recording pipette. Each point is the current during a step to -100 mV delivered from a holding potential of +30 mV. **(D)** The I-V relation measured with a 100-ms voltage ramp from - 100 to +100 mV.

### JF646-BAPTA blinks from transitions between singlet and triplet dark states

Single-channel gating events are expected to cause binary fluctuations in JF646-BAPTA fluorescence due to rapid binding of Ca^2+^ upon opening and rapid loss of Ca^2+^ by diffusion after channel closure. However, such events could be easily conflated with reversible photodynamic transitions of the dye between the fluorescent and non-fluorescent states, a process known as blinking (Ha and Tinnefeld, 2012). The parent dye, JF646, is known to blink under reducing conditions (Grimm et al., 2015; Guthrie et al., 2020), and this feature has been enhanced and exploited in modified versions to generate superresolution images (Holland et al., 2024). However, blinking of JF646-BAPTA has not been reported to date.

To test for intrinsic blinking behavior of JF646-BAPTA, we measured single-molecule fluorescence under conditions of complete Ca^2+^ saturation. Cells expressing mCh-STIM1 and labeled Orai1-HaloTag were store depleted as described in Fig. 1, and treated with digitonin to permeabilize the plasma membrane in the presence of saturating Ca^2+^ (10 mM Ca^2+^). Under these conditions, strong laser excitation (15 mW) caused stepwise bleaching over 30-60 s (**Fig. 2A**). In nearly every case, bleaching steps were accompanied by transient increases in fluorescence, a hallmark of blinking. These prominent fluctuations were most easily seen after the fluorescence had declined to the level of a single remaining JF646-BAPTA (**Fig. 2A, B**). Because of saturating Ca^2+^ conditions and the permeabilized PM, these fluctuations cannot be the result of channel gating events; instead, their amplitude is consistent with dye blinking. The dye signal transitions to a level that is indistinguishable from the local background measured adjacent to the STIM/Orai punctum and therefore represents a completely non-fluorescent state (**Fig. 2A**, expanded trace). The dwell times of single JF646-BAPTA molecules in the bright and dark states followed double exponential distributions with time constants of 89 and 609 ms for the bright state and 72 ms and 1166 ms for the dark state (**Fig. 2B**). In addition, this rapid blinking behavior was sometimes observed following long bright or dark periods lasting seconds to tens of seconds (**Fig. 2A**, right). The transitions between the slower and more rapid blinking kinetics may be related to slow changes in the dye environment, but the underlying mechanisms were not pursued further.

**Figure 2.**
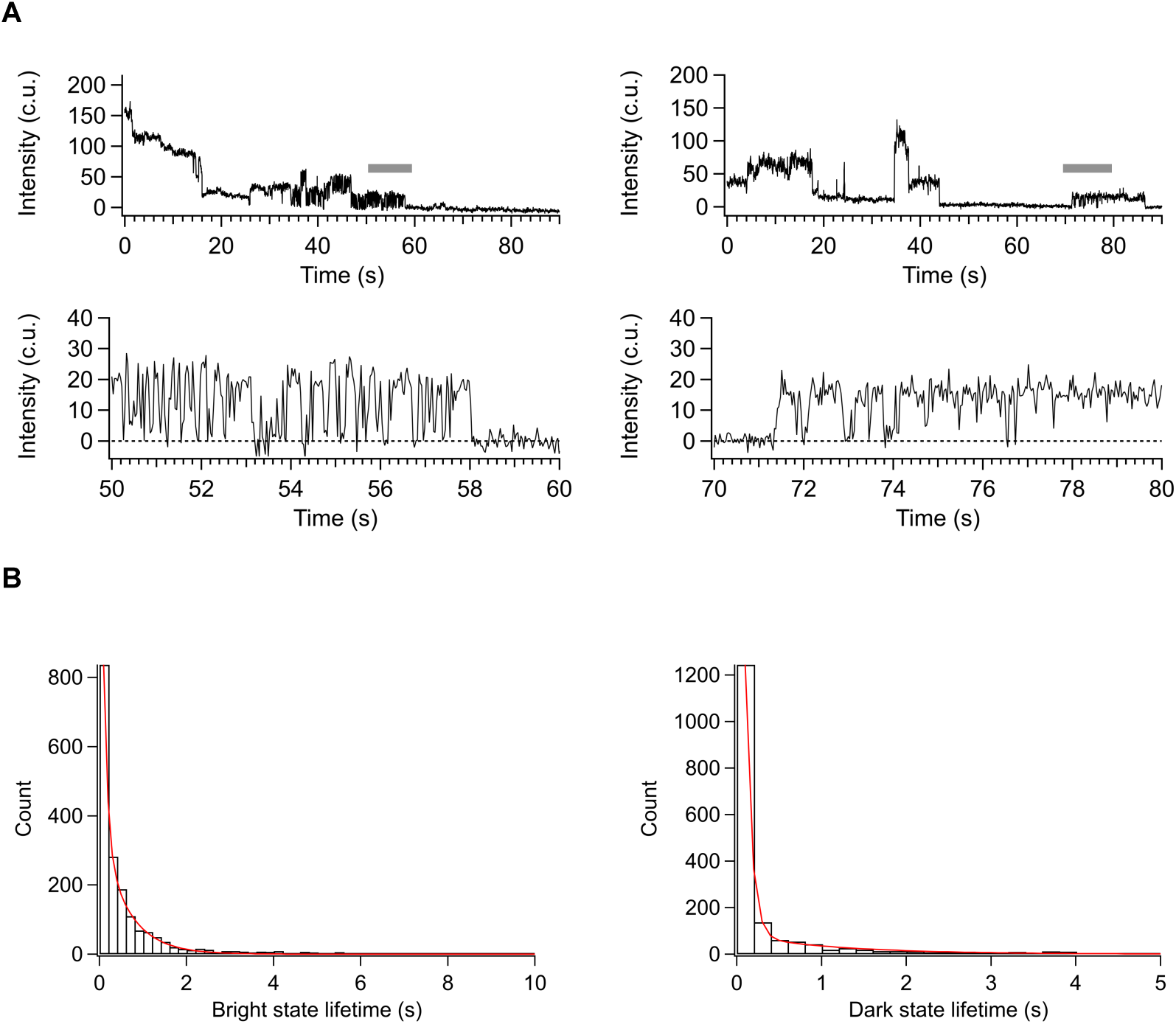
Orai1-HaloTag labeled with JF646-BAPTA exhibits blinking in saturating Ca^2+^. COS-7 cells expressing Orai1-HaloTag and mCh-STIM1 were treated with TG to induce puncta formation before exposure to 20 µM digitonin and 10 mM Ca^2+^ to saturate JF646-BAPTA. **(A)** *Left*, after stepwise bleaching of one channel to a single active JF646-BAPTA dye, rapid fluctuations appear. *Right*, spontaneous recovery of fluorescence in a second channel after a prolonged dark state period. The gray bars mark sections expanded below to show transitions to the zero-fluorescence level. 33 ms exposure with 3-frame boxcar averaging, 15 mW laser power. **(B)** Dwell time histograms of the bright and dark events like those shown in A. A biexponential fit to the bright state distribution (τ_1_=89 ms, A_1_=0.62; τ_2_=609 ms, A_2_=0.38) and biexponential fit to the dark state distribution (τ_1_=72 ms, A_1_=0.94; τ_2_=1166 ms, A_2_=0.06) are shown. Data compiled from 99 channels from 2 cells. In this and all subsequent figures, c.u. corresponds to camera output units, and unless otherwise noted, all values have been corrected for local background fluorescence (see Methods).

Because fluorescent dye blinking is known to be affected by multiple factors including the redox environment and pH (Ha and Tinnefeld, 2012), we also examined dye behavior in intact cells. In store-depleted cells where we would expect to see fluorescence fluctuations due to channel gating, we observed clear fluctuations after the laser had bleached all but one of the JF646-BAPTA ligands in a punctum (**Fig. 3A**). While these might at first appear to represent gating events, their characteristics suggest they are due instead to dye blinking. As shown in **Fig. 3A**, the signals in intact cells transition to and from the background level (indicating a completely non-fluorescent state), whereas a channel closing event would return the local [Ca^2+^]_i_ to a resting level of ∼100 nM, a level expected to produce at least 20% of the saturated state intensity based on the dynamic range of JF646-BAPTA in vitro (Deo et al., 2019) and ∼50% of the saturated state intensity based on in situ measurements (described below). Moreover, the transitions to the dark level are complete within a single 10-ms exposure (**Fig. 3B**), faster than expected from the Ca^2+^ unbinding rates of BAPTA (Naraghi, 1997), fura-2 (Kao and Tsien, 1988), or JF646-BAPTA (τ=10-30 ms, see below). Blinking events in intact cells had similar kinetics to events observed under Ca^2+^-saturated conditions, with time constants of 217 and 1030 ms for the bright state, and 156 and 650 ms for the dark state (**Fig. 3C**). Additional very long bright and dark states lasting tens of seconds were also apparent, similar to those we observed under saturated conditions (**Fig. 3A, B**). Together, these measurements show that JF646-BAPTA blinks in intact cells at physiological [Ca^2+^]_i_, making it difficult or impossible to identify and quantify true gating events from the fluorescence fluctuations of single dyes attached to the channel.

**Figure 3.**
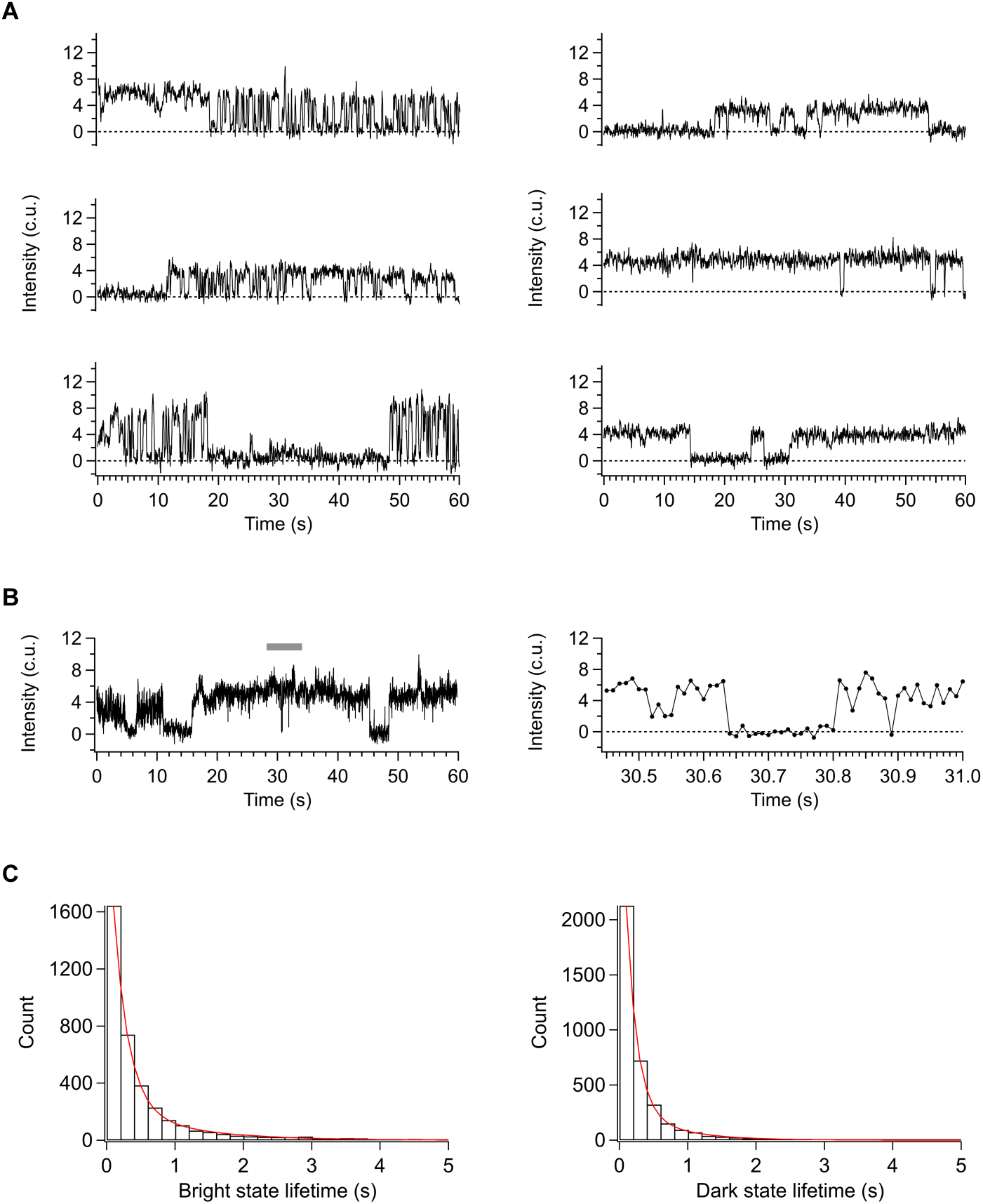
Blinking behavior of Orai1-HaloTag labeled with JF646-BAPTA in intact cells after ER Ca^2+^ depletion. COS-7 cells expressing mCh-STIM1 and Orai1-HaloTag labeled with JF646-BAPTA were treated with 2 µM CPA in the presence of 20 mM extracellular Ca^2+^. **(A)** Representative single-channel traces show fluctuations to the zero-fluorescence level alternating with long-lived bright or dark events. 20 ms exposure, 3 mW laser power, 5 frame boxcar averaging. **(B)** Transitions to and from the zero-fluorescence level are complete within the single-frame exposure time of 10 ms. The section marked by the gray bar is expanded to the right. 9 mW laser power. **(C)** Dwell time histograms of bright and dark events. Biexponential fits to the bright state distribution (τ_1_=217 ms, A_1_=0.86; τ_2_=1030 ms, A_2_=0.14) and dark state distribution (τ_1_=156 ms, A_1_=0.86; τ_2_=650 ms, A_2_=0.14) are shown. Data compiled from 128 channels from 4 cells.

### Characterizing the response properties of JF646-BAPTA bound to Orai1-HaloTag

In addition to the problems caused by blinking, recording JF646-BAPTA signals in intact cells presents other challenges to recording single-channel events. The resting potentials of HEK293 and COS-7 cells are relatively depolarized (-24 and -35 mV, respectively) (Kirkton and Bursac, 2011; Ishii et al., 1996) and may vary over time, generating a low and variable driving force for Ca^2+^ entry. Given its K_d_ of 150 nM, JF646-BAPTA can respond to increases in cell-wide global [Ca^2+^]_i_, which will contaminate the local signal produced by the attached channel and reduce the effective dynamic range of the dye; treating cells with a fixed amount of EGTA/AM to prevent these global changes generally produces a range of cytosolic buffer concentrations and because of cell-to-cell differences in Ca^2+^ influx cannot be assumed to consistently buffer Ca^2+^ over time in each cell. Finally, the speed with which Ca^2+^ influx can be reversibly stopped by extracellular perfusion with Ca^2+^-free solutions is limited by the slow rate of solution exchange in the restricted space between the coverslip and the plasma membrane [5-10 s for 90% exchange (Demuro and Parker, 2005)].

To address these challenges, we applied an approach based on a combination of whole-cell voltage clamp recording and TIRF imaging (‘Patch-TIRF’). After break-in, EGTA (12 mM) and IP_3_ (10 µM) in the pipette quickly depletes the ER Ca^2+^ store and activates Orai1 while holding [Ca^2+^]_i_ constant at low nM levels (Zweifach and Lewis, 1995), and voltage clamp provides rapid control of the driving force for Ca^2+^ entry. In the experiment of **Fig. 4**, we applied Patch-TIRF to test whether JF646-BAPTA on Orai1-HaloTag reports only Ca^2+^ influx from Orai1 channels and not from other potential Ca^2+^ sources in the neighborhood of the labeled Orai1. Brief hyperpolarizations from the holding potential of +30 mV to -100 mV evoked transient increases in JF646-BAPTA fluorescence, and La^3+^ (100 µM) blocked the response as expected for Orai1 (**Fig. 4B**). In contrast, hyperpolarization failed to evoke JF646-BAPTA signals in cells expressing the non-conducting E106A Orai1-HaloTag mutant (Prakriya et al., 2006; Yeromin et al., 2006) (**Fig. 4C**), demonstrating that under our whole-cell recording conditions, JF646-BAPTA selectively indicates Orai1 activity.

**Figure 4.**
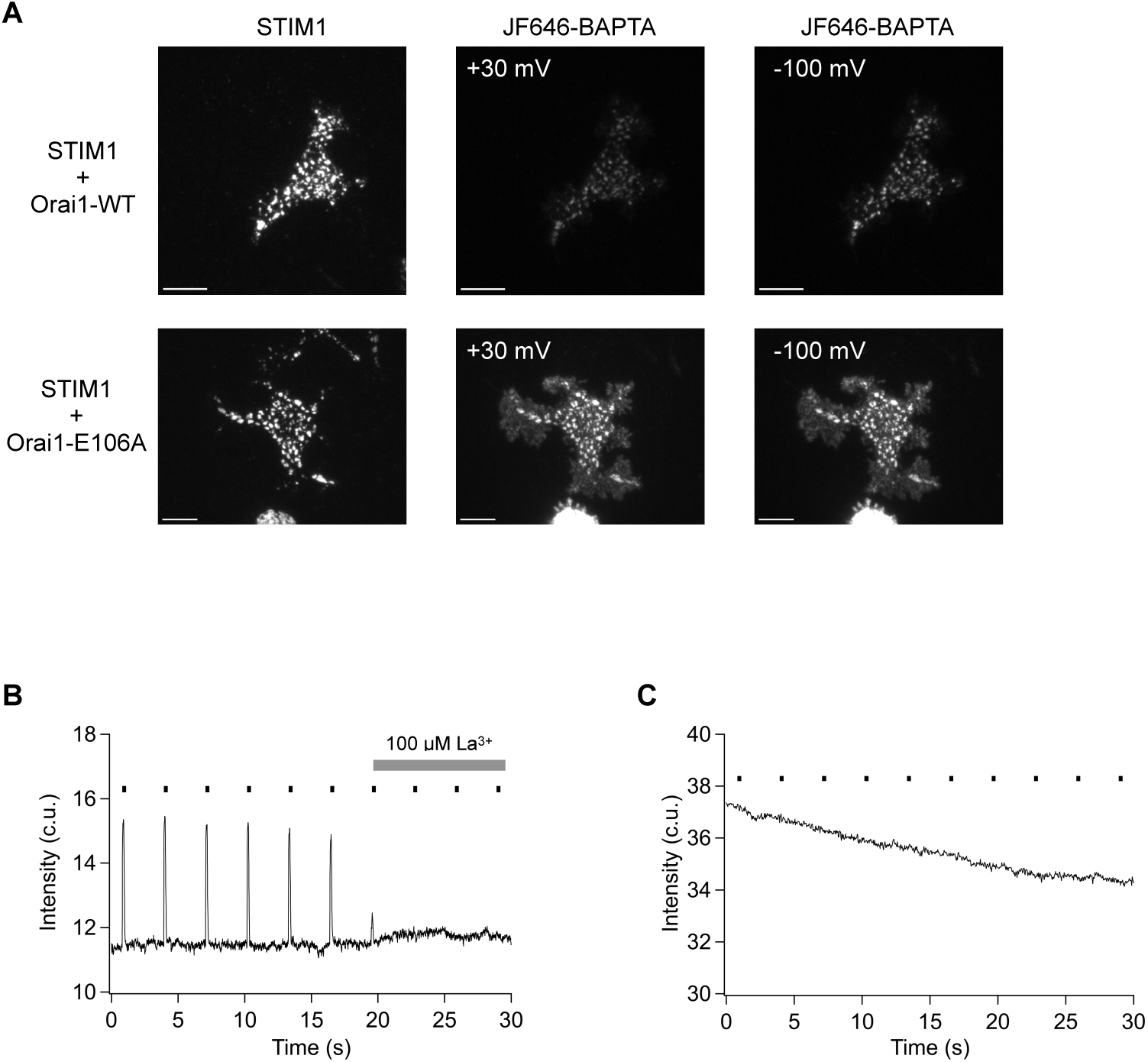
JF646-BAPTA selectively reports Ca^2+^ entering through Orai1 channels during whole-cell recording. **(A)** Images of HEK293 cells expressing mCh-STIM1 and either WT Orai1-HaloTag (*top row*) or Orai1-E106A-HaloTag (*bottom row*). The fluorescence of JF646-BAPTA increases at - 100 mV for WT Orai1 but not Orai1-E106A, despite being co-clustered with STIM1. Bar=10 µm. **(B)** Fluorescence of WT Orai1-HaloTag across the cell footprint increases during 100-ms pulses to -100 mV from a holding potential of +30 mV (bars). The response is blocked by 100 µM La^3+^ as expected for influx through Orai1. **(C)** Fluorescence of Orai1-E106A-HaloTag across the cell footprint does not respond to hyperpolarizing pulses, indicating that the fluorescence increase of WT Orai1 derives directly from Ca^2+^ flux through Orai1. Orai1-E106A-HaloTag expression level in C was ∼3-fold higher than in B in order to optimize detection of even a small amount of Ca^2+^ from other sources. 10 ms exposure, 1 mW laser power.

We applied the voltage clamp to vary the driving force for Ca^2+^ entry and find conditions that produce minimum and maximum fluorescence corresponding to the Ca^2+^-free and Ca^2+^- saturated forms of JF646-BAPTA. Repetitive 10-s voltage ramps from -100 to +100 mV evoked a series of fluorescence responses in single Orai1 puncta in the presence of 20 mM Ca^2+^ (**Fig. 5A**). Responses differed among individual puncta, showing that each punctum responds independently to store depletion. The average of the four ramp responses from 21 puncta showed the sensitivity of the JF646-BAPTA response to membrane potential (**Fig. 5B**). The fluorescence-voltage relation is consistent with the voltage dependence of Ca^2+^ current (**Fig. 1D**), with a minimum clearly greater than zero reached at potentials more positive than ∼+20 mV, and a steep increase at negative potentials. Fluorescence intensity reached a limiting value at -75 mV and below, indicating saturation of the dye near the channel pore at these rates of Ca^2+^ entry. Thus, for an open channel the intensity of JF646-BAPTA is maximized at -100 mV while the minimal fluorescence at +30 mV mimics the response of a closed channel.

**Figure 5.**
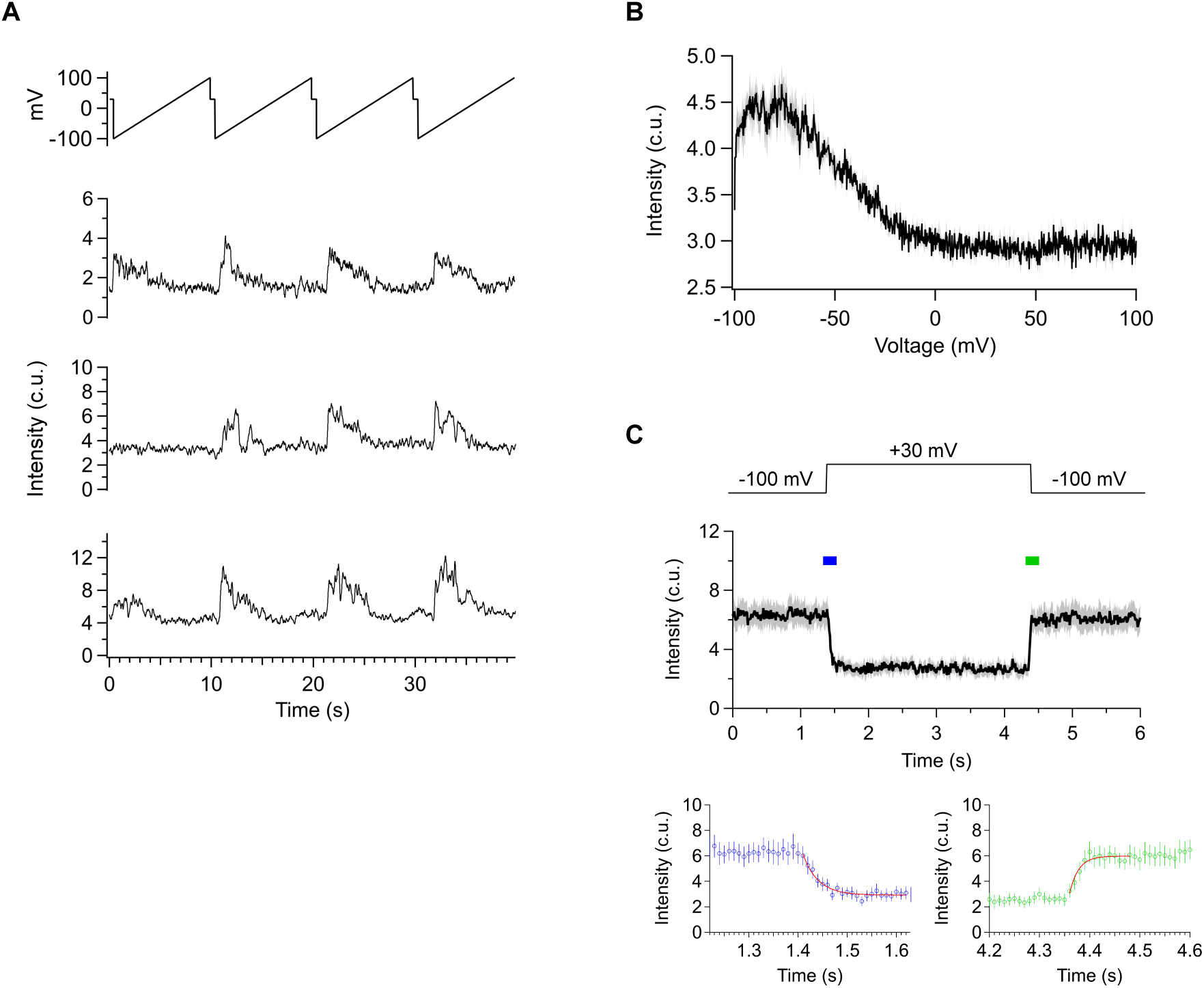
In situ characterization of Orai1-HaloTag labelled with JF646-BAPTA. HEK293 cells expressed mCh-STIM1 and Orai1-HaloTag labeled with JF646-BAPTA. All data are shown relative to local background fluorescence. **(A)** Responses of 3 puncta to voltage ramps from -100 to +100 mV. **(B)** Saturation of JF646-BAPTA in cells expressing Orai1-HaloTag. Mean fluorescence ± SEM was compiled from the responses of 22 puncta to 4 voltage ramps like those in A. Binding kinetics of Ca^2+^ for JF646-BAPTA reduces the response over the first several points of the ramp (- 100 to -99 mV). The response is minimal at potentials above ∼+20 mV, and saturates at potentials of -75 mV or below. **(C)** Kinetics of JF646-BAPTA fluorescence in response to a voltage step from +30 to -100 mV. Expanded sections indicated by colored bars show single-exponential fits to the off and on transitions (τ_off_=30 ms, τ_on_=18 ms). Mean ± SEM of 6 puncta. In A-C, 10 ms exposure, 1 mW laser power. A had 15 frame boxcar averaging.

The Ca^2+^ binding and unbinding kinetics and overall dynamic range of JF646-BAPTA in cells were estimated by rapidly changing the driving force for Ca^2+^ entry (**Fig. 5C**). Stepping the voltage from -100 mV to +30 mV terminated influx, causing puncta fluorescence to decline to a minimum due to Ca^2+^ unbinding and diffusion into the cytosol, with a single-exponential time constant of 30 ms for Ca^2+^ unbinding (τ_off_). Stepping from +30 to -100 mV restored influx and fluorescence increased to a maximum value within 18 ms (τ_on_). The on and off times are distinctly slower than the transition times for blinking (<10 ms, **Fig. 3B**). The steady-state reduction of fluorescence during the step from -100 to +30 mV indicates a maximal dynamic range of 2.4. Together, the kinetics and dynamic range of JF646-BAPTA provide useful criteria for distinguishing gating from blinking.

Multiple Orai1 channels can accumulate with STIM1 in a single punctum and are not easily distinguished if spaced closer than the diffraction limit (∼200 nm). To establish criteria for identifying single-channel puncta, we measured puncta fluorescence intensity from cells expressing mCh-STIM1 and Orai1-HaloTag. We adopted a standard set of recording conditions (1 mW laser power, 33 ms exposure, 3-frame boxcar averaging) that allowed extended imaging over 60 s without significant bleaching while maintaining an adequate signal-to-noise ratio (SNR). An example of Orai1-JF646-BAPTA puncta from one cell at -100 mV shows a range of fluorescence intensities among individual puncta (**Fig. 6A**). The fluorescence due specifically to Ca^2+^ influx at each punctum was quantified as the increase in fluorescence when the membrane was hyperpolarized from +30 mV to -100 mV (**Fig. 6B**). In measurements from 290 puncta, the intensity distribution displayed multiple peaks (**Fig. 6C**). The lowest two peaks were centered at 5.1 and 11 c.u., which we interpret as the response amplitudes of 1 and 2 channels, respectively. This assignment of the single-channel intensity allows a rough estimate of the number of Ca^2+^-sensitive dyes per channel. A single JF646-BAPTA generates a signal of 13.5 ± 1.1 c.u. (mean ± sem; n=17), measured from bleaching steps under Ca^2+^ -saturated conditions at 15 mW laser power in the experiment of **Fig. 2A**. This is equivalent to 0.9 c.u. under our standard imaging conditions (1 mW laser power), suggesting that each channel has on average 5.1/0.9, or ∼6 Ca^2+^-responsive dyes, consistent with complete labeling by JF646-BAPTA (**Fig. S1**). The observed spread of the single-channel intensity distribution potentially has multiple sources, including variation in the number of responsive dyes, degree of blinking, non-uniform TIRF laser illumination across the imaging field, and variations in the Z-position of the plasma membrane within the exponentially decaying evanescent field (Fish, 2022).

**Figure 6.**
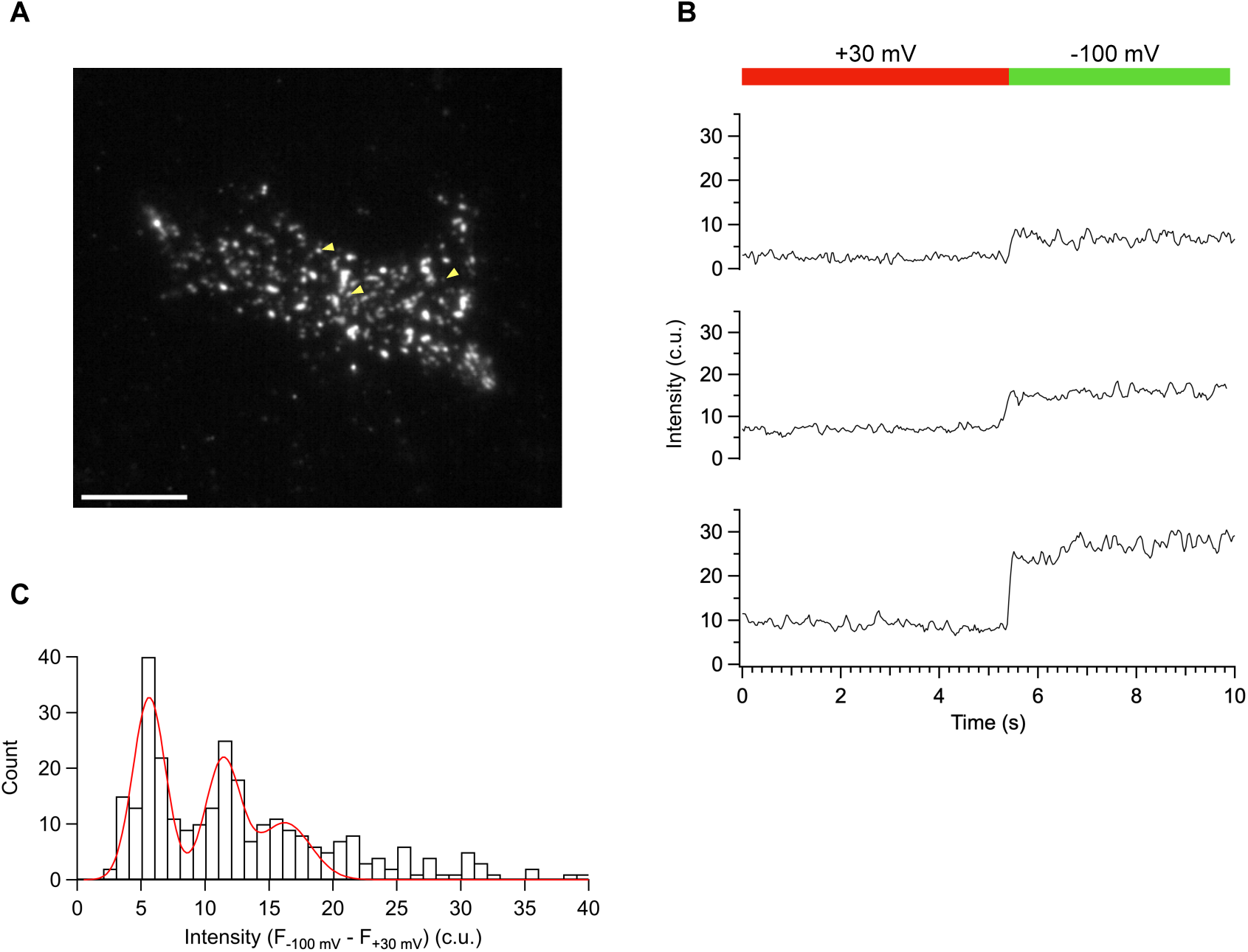
Identification of puncta containing a single Orai1-HaloTag channel. **(A)** TIRF image of Orai1-HaloTag in a HEK293 cell coexpressing mCh-STIM1 during whole-cell recording at -100 mV. Note the range of JF646-BAPTA fluorescence intensities among different puncta. Arrowheads mark puncta plotted in B. Bar=10 µm. **(B)** Traces of single Orai1-HaloTag puncta from A upon hyperpolarization to -100 mV from a +30 mV holding potential, showing examples of 1-, 2-, and 3-channel puncta. **(C)** Distribution of fluorescence intensities at -100 mV relative to the baseline fluorescence at +30 mV, compiled from 289 puncta in 10 cells. Gaussian fits to the histogram indicate peaks at 5.1, 10.9, and 15.8 c.u., corresponding to 1-, 2-, and 3-channel puncta. In B and C, 33 ms exposure, 1 mW laser power, 2 frame boxcar averaging.

### Gating behavior of single Orai1 channels measured with JF646-BAPTA

The characterization of JF646-BAPTA properties and Orai1-HaloTag labeling enabled us to develop criteria for identifying gating events and assessing the kinetic behavior and open probability of single Orai1 channels after store depletion. Whole-cell recording was initiated at a holding potential of +30 mV, and image acquisition was started ∼30 s later to allow time for store depletion and full activation of Orai1 channels. After recording baseline fluorescence at +30 mV to define the closed channel fluorescence level, the holding potential was hyperpolarized to -100 mV. Orai1-JF646-BAPTA puncta were selected for analysis based on several criteria: colocalization with GFP-STIM1, a stable fluorescence baseline at +30 mV which increased by 3-6 c.u. at -100 mV (typical for a single channel; **Fig. 6C**), and a signal-to-noise ratio (SNR) at -100 mV of >4 (see Methods).

Figure 7A shows recordings from several Orai1-HaloTag channels that met these selection criteria. In each case, fluorescence increased rapidly upon hyperpolarization and after periods of 5-15 s showed stepwise fluctuations back to the baseline (+30 mV) level, which we determined to be the dye fluorescence in the absence of influx (Fig. 5B). These transitions satisfy the criteria for a closing event, and confirm that the signals arise from a single channel, as it is unlikely that multiple independently gated channels would close synchronously during such prolonged openings.

**Figure 7.**
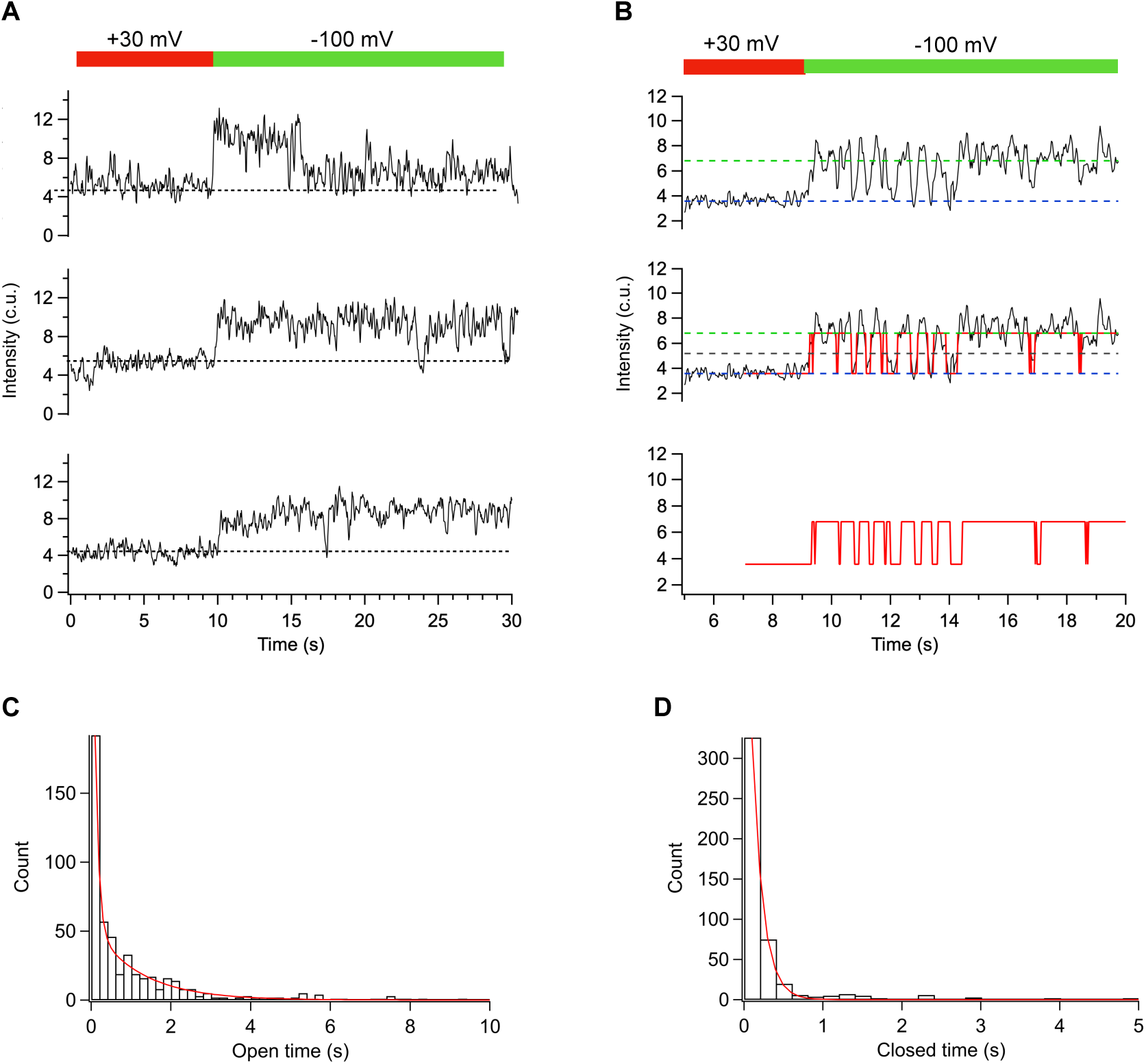
Single-channel behavior of Orai1-HaloTag during whole-cell recording. HEK293 cells expressed mCh-STIM1 and Orai1-HaloTag labeled with JF646-BAPTA. All data are shown relative to local background fluorescence. **(A)** Examples of single-channel responses to hyperpolarization to -100 mV. In each case, fluorescence is ∼5 c.u. above the 0-current level (defined as the intensity at +30 mV, dashed line) and drops to the 0-current level during closing events. **(B)** Thresholding procedure for detecting gating transitions. Levels were set for the open and closed states from fluorescence intensities at -100 and +30 mV, respectively (*top*). A 50% threshold was applied to detect transitions between open and closed states (*middle*), to generate an idealized trace (*bottom*). **(C)** Open dwelltimes follow a biexponential distribution (τ_1_=0.09 s, A_1_=0.74; τ_2_=1.19 s, A_2_=0.26). **(D)** Closed dwelltimes were exponentially distributed (τ=0.14 s). Data compiled from 36 channels in 12 cells. In A-D, 33 ms exposure, 1 mW laser power, 3 frame boxcar averaging.

To measure open and shut durations, we fit the closed and open state amplitudes based on the fluorescence at +30 mV and -100 mV, respectively, and identified transitions between states as crossing a 50% threshold (Fig. 7B). Each trace yielded an idealized waveform for measuring open and closed dwell times. From 36 traces that met our criteria for analysis (**Fig. S3**), open dwell times were biexponentially distributed with time constants of 92 ms (A_1_=74% of events) and 1190 ms (A_2_=26% of events) (Fig. 7C). Closed dwell times were exponentially distributed with a time constant of 140 ms (Fig. 7D). The overall proportion of time spent in short and long events, calculated from A x τ for each, was 18% for short openings and 82% for long openings.

Much longer openings and closings lasting tens of seconds were also detected but were too sparse to be fitted accurately as components of the dwell time distributions. Long events also tended to begin before the time of hyperpolarization or continue beyond the end of the recording period (e.g., see Fig. 7A), precluding measurement of an open or closed duration. In particular, of 559 single-channel puncta in 3 cells, 11% of channels appeared to fully lack activity over a recording time of 30 s despite colocalizing with mCh-STIM1 (Fig. 8A, B). These ‘silent’ channels would be undetectable by electrophysiological assays but were identifiable by fluorescence levels of 2-3 c.u. above the local background that did not increase with hyperpolarization, consistent with Ca^2+^-free single channels. Combining these data with open probabilities measured from 28 active channels yields a skewed distribution, with the majority of channels exhibiting a high P_O_ ≥ 0.7 (Fig. 8C).

**Figure 8.**
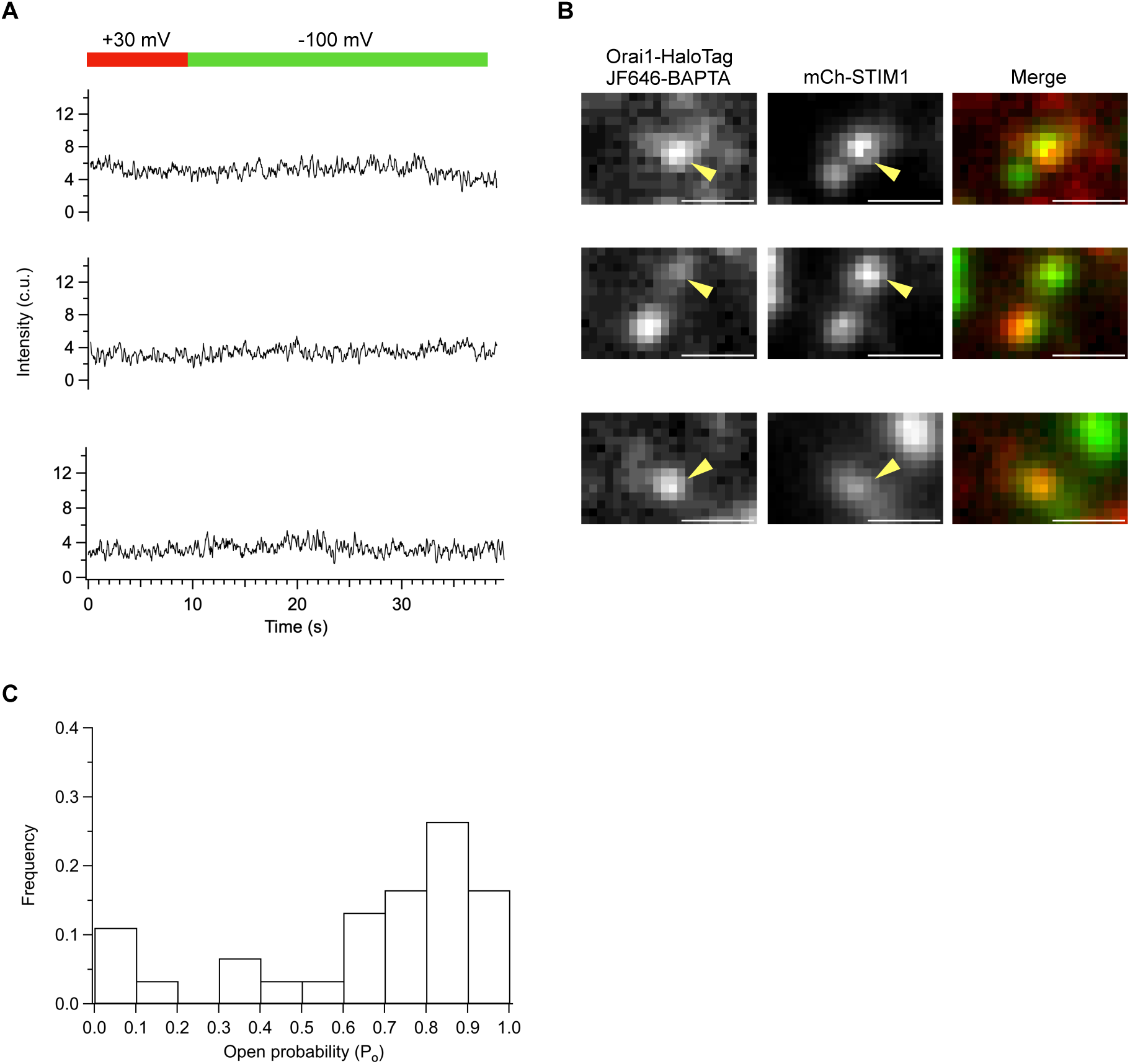
Open probability of Orai1-HaloTag channels during whole-cell recording. HEK293 cells expressed mCh-STIM1 and Orai1-HaloTag labeled with JF646-BAPTA. **(A)** Example traces showing single channels with no apparent activity over a period of 40 s (‘silent channels’). **(B)** TIRF images of Orai1-HaloTag with arrowheads showing the channels displayed in A with merged images showing colocalization of mCh-STIM1 (green) and Orai1-HaloTag (red). Bar=1 µm. **(C)** The relative frequency of P_O_ values from ‘silent’ channels (out of 559 channels at STIM1 puncta) and 28 active channels from Fig. 7C and D.

## DISCUSSION

Understanding the kinetic behavior and gating mechanisms of the CRAC channel has lagged behind that of many other channels because a characteristic feature – an extremely small 20-40 fS unitary conductance – precludes single-channel recording. Although optical monitoring of local Ca^2+^ with JF646-BAPTA bound to Orai1-HaloTag offers a potential solution to this conundrum, we found that reversible blinking to a non-fluorescent state presents a serious challenge to the detection of true gating events. However, by analyzing the response of multiple dyes bound to each channel, we were able to identify channel openings and closings of single CRAC channels in response to Ca^2+^ store depletion. These results provide a new level of mechanistic detail about CRAC channel gating that could not have been gained from macroscopic recordings and offer an approach for further dissecting gating mechanisms, while revealing important caveats for the use of Ca^2+^ sensitive dyes like JF646-BAPTA to monitor single-channel behavior more generally. Protein indicators (GECIs) may present similar complications for single-channel analysis, as GCaMP8 fused to Orai1 has been shown to undergo reversible photoinactivation, probably due to cis-trans photoswitching of the fluorophore (Dynes et al., 2023).

### Blinking behavior of JF646-BAPTA can resemble single-channel events

JF646-BAPTA offers advantages for Ca^2+^ measurements in cells but not without limitations. For monitoring Ca^2+^ in small volumes, JF646-BAPTA performs better than currently available Ca^2+^- sensitive GECIs, with greater brightness and photostability, as well as faster Ca^2+^ binding and unbinding kinetics (Deo et al., 2019). Its high sensitivity in situ has been demonstrated in Ca^2+^ measurements within femtoliter compartments like primary cilia (Deo et al., 2019) and as a spatial reporter of PIEZO1 channel activity in several cell types and tissue organoids derived from human induced pluripotent stem cells (Bertaccini et al., 2025). However, analysis of single-channel activity is complicated by the tendency of JF646-BAPTA to blink. Two hallmarks of blinking distinguish this behavior from gating events: transitions to a completely non-fluorescent state and transition kinetics that are faster than the Ca^2+^ unbinding kinetics of JF646-BAPTA. In a recent study of PIEZO1-HaloTag labeled with JF646-BAPTA (Bertaccini et al., 2025), signals fluctuated from a well-resolved bright level to a zero-fluorescence level rather than the finite fluorescence expected for Ca^2+^-free dye attached to HaloTag in vitro (∼20% of maximum) (Deo et al., 2019) or to Orai1-HaloTag in situ (∼50% of maximum; Fig. 5C). While these events were interpreted as single-channel openings, their characteristics are consistent with blinking of a single JF646-BAPTA attached to the channel rather than gating transitions and suggest that additional tests are needed to distinguish single-channel gating events from photophysical dye behavior.

Dye blinking can occur through several mechanisms which produce distinct kinetic signatures. These include entry into a non-fluorescent triplet state, redox effects, protonation of the fluorophore, cyclization of the dye, and electromagnetic interaction with nearby fluors (Di Fiori and Meller, 2010; Ha and Tinnefeld, 2012). We do not know the source(s) of blinking in the case of JF646-BAPTA, but the existence of multiple bright and dark kinetic states (Fig. 3) suggests that more than one mechanism may be involved.

### Patch-TIRF recording can be used to distinguish gating from blinks

Varying the driving force for Ca^2+^ entry with voltage ramps indicated the minimum and maximal signals obtainable from JF646-BAPTA in situ (Fig. 5). The fluorescence was constant and minimal at voltages between ∼0 and +100 mV; at these voltages inward I_CRAC_ is negligible, and the 12 mM EGTA in the pipette is expected to buffer the free [Ca^2+^]_i_ to low nM levels. This fluorescence represents the level expected when the channel is closed, and unlike the dark state is well above zero. At voltages below -75 mV, fluorescence reached a maximum, representing the maximal signal possible from Ca^2+^-saturated JF646-BAPTA. The response to steps from +30 to -100 mV indicated a maximum dynamic range for JF646-BAPTA of 2.4, which is lower than the reported range of ∼5 reported for JF646-BAPTA bound to free HaloTag in vitro (Deo et al., 2019). There are several possible explanations for this discrepancy. One is that the quenching of the Ca^2+^-free dye or dequenching upon Ca^2+^ binding may be reduced by proximity effects within the cluster of HaloTags attached to the channel. Another is that JF646-BAPTA/AM may be incompletely de-esterified in the cell, perhaps as a consequence of poor accessibility of esterases to the dye after it is bound to the Orai1-HaloTag; a fraction of Ca^2+^-independent fluorescence would be expected to reduce the dynamic range. Regardless of the cause, the effective dynamic range may be specific to the local environment of the dye and should therefore be assessed directly, e.g., using Patch-TIRF, for each application to a specific channel or cell type.

To capture long duration events, we applied a low laser intensity (1 mW) that minimized bleaching over extended imaging times. While this enabled detection of very long events (>50 s), the signals were small and susceptible to fluctuations contributed by shot noise, dye blinking, and movement of the plasma membrane in the Z-axis during TIRF imaging. The sampling rate (33 ms with an additional 3-frame boxcar average) that was required to achieve a usable SNR causes us to miss events shorter than ∼100 ms. Conversely, high frequency noise may increase the number of apparent short openings or closings, while interrupting long closings or openings and leading to underestimates of closed and open times. These limitations could be reduced with higher laser intensity to improve the SNR and obtain more detailed kinetic information over shorter durations. Most importantly, development of Ca^2+^-sensitive HaloTag dyes that are less prone to blinking would greatly improve the SNR and simplify the analysis of single-channel events.

### The kinetic behavior of single Orai1 channels

Single-channel gating events were distinguished from blinking by rapid transitions between a defined open-channel fluorescence intensity and the 0-current level (indicated by the intensity at +30 mV). Kinetic analysis of the single channel responses reveals the behavior of CRAC channels activated by STIM1 under conditions of full store depletion. Open time distributions indicate mean open times of 0.09 and 1.19 s and a mean closed time of 0.140 s, with longer events lasting tens of seconds that could not be adequately sampled within the achievable recording time.

The shorter STIM1-evoked openings display similar kinetics to single-channel events first described by Dynes et al. for G-GECO1-Orai1 in the presence of low concentrations of CAD, which consisted of ‘flicker’ events lasting several hundred ms and step-like ‘pulses’ lasting from one to several s (Dynes et al., 2016). Longer openings were not reported in that study, perhaps because Orai1 was activated by a low concentration of soluble STIM1 CAD fragment, which is likely to bind less tightly than ER-anchored STIM1 to Orai1 and only infrequently reach a level of binding necessary to open the channel. The step-like events we observed are clearly distinct from the ∼1.5-s oscillatory responses of G-GECO1.2-Orai1 that were reported in response to high CAD expression or to STIM1 in store-depleted cells (Dynes et al., 2016). While the reasons for this discrepancy are not clear, several experimental factors in the previous study may have contributed, such as the slower kinetics of G-GECO1.2 and sampling rate (100 ms exposure and 4-frame averaging), the lack of voltage control, and crosstalk of signals within channel clusters enabled by the absence of exogenous intracellular Ca^2+^ buffering.

The long lifetimes of CRAC channel openings we observed (seconds to tens of seconds) may have important implications for the functions of SOCE. From the events we could capture in the recording time of 30-60 s, the majority of open time (82%) was spent in the 1.19-s mean duration class, and even longer openings (tens of s) were apparent but could not be adequately sampled given the maximum recording time of 30-60 s. Seconds-long openings may enhance the specificity of Ca^2+^ signaling through Orai1, by enabling the few low-conductance Orai1 channels present at each ER-PM junction to effectively promote local processes such as transcription through calcineurin activation and NFAT dephosphorylation (Kar et al., 2021) and store refilling by SERCA that permits Ca^2+^ tunneling through the ER to activate specific targets (Courjaret et al., 2025), all while minimizing Ca^2+^ spillover into the cytosol.

Interestingly, a substantial proportion (11%) of Orai1-HaloTag channels localized at STIM1 puncta did not appear to conduct Ca^2+^ over recording periods of ∼30 s. A population of such ‘silent’ channels may help explain previous observations in a study of Orai1 channels labeled with GCaMP6f, where the measured local [Ca^2+^]_i_ at puncta was much lower than expected based on estimates of the number of Orai1 channels and their open probability (Dynes et al., 2020). Inactive Orai1 channels may be a reserve that is recruited by specific stimuli or conditions and could reside in the plasma membrane or in recycling endosomes docked at ER-PM junctions (Hodeify et al., 2015). Optical channel recording offers a unique approach to specifically identify and study this population of inactive, electrically silent channels.

A previous study using non-stationary fluctuation analysis to evaluate changes in P_O_ during whole-cell CRAC current activation and deactivation concluded that upon store depletion, CRAC current activates by recruitment of ‘silent’ channels to a highly active (P_O_ ∼0.8) state (Prakriya and Lewis, 2006). Our single-channel measurements support this notion, as shown by the distribution of P_O_ values with the majority of channels having a high P_O_ ≥0.7. A limitation of this noise analysis approach is that it was based on a series of 200-ms current recordings, which would miss long closing closures and hence overestimate the overall P_O_ (Yen and Lewis, 2018). Optical recording offers for the first time the ability to measure the true P_O_ by enabling long continuous recordings that can directly capture slow gating events. Optical recording may be applied in the future to explore the structural basis for long channel openings, including a potential role of an M101-F99 ‘latch’ implicated in stabilizing the open state (Yeung et al., 2020), as well as the nonlinear effects of STIM:Orai binding stoichiometry on activity (Yen and Lewis, 2018). In addition, CRISPR-mediated HaloTag labeling of endogenous Orai channels will allow gating to be studied in native settings, and ultimately follow single CRAC channels as they encounter STIM1 at ER-PM junctions in response to physiological levels of store depletion. More broadly, HaloTag-based optical recording offers a powerful approach to study a variety of Ca^2+^-permeable channels at single-channel resolution across large expanses of the cell, while offering access to restricted compartments like neuronal and immune synapses and primary cilia, where small numbers of channels exert an outsized influence on cell physiology.

## METHODS

### Cell culture

Orai1/2/3 triple-knockout HEK293 cells (Yoast et al., 2020) and COS-7 cells (ATCC) were maintained in in high-glucose DMEM supplemented with 10% FBS and 1% penicillin/streptomycin at 37° C with 5% CO_2_. Throughout this study any reference to ‘HEK293 cells’ refers directly to the triple-knockout HEK293 cells. All cells were propagated at 70-80% confluency in 6-cm dishes coated with 1% Matrigel (20 min at 37° C; Corning) using TrypLE (Gibco).

### Plasmid construction

Orai1-HaloTag was generated from the Orai1-GCaMP6f construct (Addgene) by replacing GCaMP6f with HaloTag (Promega). The HaloTag insert was amplified by PCR using the forward primer 5ʹ-GCGGGCCCGGGATCCACCGATGGCAGAAATCGGTACTGGC-3ʹ and reverse primer 5ʹ-ATGATCTAGAGTCGCGGCCGCTTTAGCCGGAAATCTCGAGCGT-3ʹ. The resulting fragment and vector backbone were digested with *BamHI* and *NotI*, followed by ligation using T4 DNA ligase (New England Biolabs). This strategy connected HaloTag to the C-terminus of Orai1 via the RILQSTVPRARDPP linker derived from the multiple cloning site used in prior Orai1-GECI studies (Dynes et al., 2016). Orai1-HaloTag-E106A was generated from Orai1-HaloTag by site-directed mutagenesis (Quikchange XL; Stratagene) using the forward primer 5’-CATGGTGGCAATGGTGGCGGTGCAGCTGGACGCTG-3’ and reverse primer 5’-CAGCGTCCAGCTGCACCGCCACCATTGCCACCATG-3’. The plasmids encoding mCherry-STIM1 and GFP-STIM1 were described previously (Wu et al., 2006). A thymidine kinase promoter–driven Orai1-HaloTag (TK-Orai1-HaloTag) construct was designed and synthesized by Twist Bioscience, and subsequently cloned into a plasmid backbone using Gibson assembly. All plasmids were validated by sequencing.

### Transfections

To assess labeling efficiency, COS-7 cells at 70-80% confluency in 6-cm dishes were transfected with 600 ng GFP-STIM1 and 600 ng TK-Orai1-HaloTag using 6 µL of Lipofectamine 3000 (Invitrogen) for 1 h. The thymidine kinase (TK) promoter did not support detectable expression of Orai1-HaloTag in HEK293 cells; therefore, for all HEK293 experiments, 600 ng mCherry-STIM1 or GFP-STIM1 and 35 ng Orai1-HaloTag driven via a CMV promoter were transfected using 6 µL of Lipofectamine 3000 for 1 h to achieve low-level expression of Orai1-HaloTag. All cells were assayed within 24 h of transfection.

### HaloTag labeling protocol

A 1 mM stock solution of JF646-BAPTA/AM in DMSO was aliquoted and stored at -20° C. For assessment of labeling efficiency several conditions were tested. 500 pM JF646-BAPTA/AM in medium for 15 min at 37° C (Bertaccini et al., 2025) produced no detectable labeling. For the protocol of Deo et al. (Deo et al., 2019), transfected cells were labeled overnight (18 h) in medium containing 1 μM JF646-BAPTA/AM at 37° C with 5% CO_2_, then washed 4 times with culture medium and incubated with 100 nM JF552 for 10 min in medium at 37° C with 5% CO_2_ before another 4x wash and 10 min recovery in the incubator before imaging. Labeling with this protocol was incomplete (**Fig. S1B**). We adopted a modified protocol to achieve full labeling, in which cells were incubated in 1 µM JF646-BAPTA/AM with 0.02% Pluronic F127 (Sigma-Aldrich) in culture medium for 1 h at 37° C with 5% CO_2_, then washed 4 times with pre-warmed medium and allowed to rest for 2 h at 37° C with 5% CO_2_ before imaging. Cells were assessed for labeling efficiency with JF552 as described above (**Fig. S1B**).

### TIRF microscopy

No. 1.5 coverglass was cleaned by sequential sonication in 100% acetone (15 min), Milli-Q water (5 min), 1 M KOH (15 min), Milli-Q water (5 min), and 100% ethanol (15 min), followed by flame treatment. Cleaned coverglass was sealed onto homemade slotted chambers using vacuum grease, and 1% Matrigel (Corning) was added for 20 min before plating labeled Orai1/2/3 triple-knockout HEK293 cells expressing GFP-STIM1 and Orai1-JF646-BAPTA in medium. After ∼4 h, the medium was exchanged with external solution containing (in mM): 130 NaCl, 20 CaCl_2_, 4.5 KCl, 1 MgCl_2_, 10 D-glucose, and 10 HEPES, adjusted to pH 7.4 with NaOH. For all experiments, we studied cells with a low level of Orai1-HaloTag expression based on the resting level of JF646-BAPTA fluorescence observed with 640 nm illumination. All experiments were performed at 22° C.

For digitonin-permeabilization experiments (Fig. 2), COS-7 cells were perfused with an external solution containing 135 mM NaCl, 10 mM CaCl_2_, 4.5 mM KCl, 10 mM D-glucose, 1 mM MgCl_2_, 10 mM HEPES, and 1 μM thapsigargin (TG) for 10 min to deplete intracellular Ca^2+^ stores. TG was then removed and replaced with 10 μM digitonin, and cells were imaged for up to 3 min under permeabilized conditions. Beyond this time window, Orai1-HaloTag puncta exhibited increased mobility.

For intact-cell experiments (Fig. 3), COS-7 cells co-expressing mCherry–STIM1 and Orai1– HaloTag were plated on Matrigel-coated coverslips on the day of imaging. Cells were incubated with 10 μM EGTA-AM in culture medium for 1 h, followed by a wash in fresh medium. After a recovery period of 1 hr, cells were exposed to 2 μM cyclopiazonic acid (CPA) in external solution containing 135 mM NaCl, 20 mM CaCl_2_, 4.5 mM KCl, 10 mM D-glucose, 1 mM MgCl_2_, 10 mM HEPES for ∼120 s prior to imaging. In some cases, intracellular Ca^2+^ stores appeared partially depleted even in the absence of CPA, as evidenced by the presence of persistent STIM1 puncta for several hours.

TIRF imaging was performed with a Nikon Ti2-E inverted microscope equipped with a TIRF-E module, Perfect Focus, and a Nikon Apo TIRF 100X Oil NA 1.49 objective. Laser illumination was provided by 488, 561, and 640 nm lasers in a LUNF laser launch (Nikon), and laser intensity was measured at the back focal plane. The following lasers and emission filters were used in series with a quad band dichroic filter (Chroma 405/488/561/638) to image GFP-STIM1 (488 nm, Chroma ET525/50m), mCh-STIM1 (561 nm, Chroma ET600/50m), and JF646-BAPTA (640 nm, Chroma ET700/75m). Images were collected with a sCMOS camera (Prime 95B, Photometrics) in 12-bit high sensitivity mode. Nikon Elements software was used to control excitation and image acquisition, and to select subregions of the field for high-speed acquisition of time series without gaps at 10 ms or 33 ms. Images of mCh-STIM1 were acquired with a single 100-ms exposure after the end of the JF646-BAPTA imaging run. Spatial registration of the GFP and JF646-BAPTA channels was done with 100-nm TetraSpeck^TM^ beads (Thermo Fisher Scientific). Line scans of beads were fit with Gaussian profiles to determine center position, and differences between the two channels were used to bring the images into alignment.

Fluorescence intensity is expressed in camera output units (c.u.). Applying the Prime 95B conversion factor of 0.5 photoelectrons/c.u. and a quantum efficiency of 0.9 photoelectrons/photon at 640 nm, 1 c.u. is equivalent to 0.56 photons.

### Whole-cell recording

Recording pipettes were pulled from borosilicate glass using a P-87 puller (Sutter Instruments) and fire-polished to achieve resistances of 2-3 MΩ. External solution contained (in mM): 130 NaCl, 20 CaCl_2_, 4.5 KCl, 1 MgCl_2_, 10 D-glucose, and 10 HEPES, adjusted to pH 7.4 with NaOH. Internal solution contained (in mM): 115 Cs aspartate, 8 MgCl_2_, 12 EGTA, 10 HEPES, and 2 Na_2_ATP, adjusted to pH 7.2 with CsOH. In all experiments except Fig. 1, 10 µM IP_3_ was added to the internal solution. Whole-cell recording was performed with an Axopatch 200B amplifier (Axon Instruments) and an ITC-16 interface (Instrutech) controlled using in-house routines in Igor Pro 6 (Wavemetrics). Command voltage was corrected for pipette junction potential. In Fig. 1C and D, a step-ramp stimulus (100 ms at -100 mV followed by a 100-ms ramp from -100 to +100 mV) was delivered every 2-3 s from a holding potential of +30 mV. Current was sampled every 200 μsec and filtered at 1 kHz and corrected for leak current recorded in the presence of 100 µM LaCl_3_ added to the external solution.

### Image analysis

Images were analyzed using ImageJ/FIJI. The TrackMate plugin was used to identify STIM1 puncta from a single image, and after verifying that all puncta had been identified, the resulting ROIs were applied to the time series of Orai1-HaloTag images. Due to some movement of Orai1-HaloTag within STIM1 puncta, every ROI was manually inspected to confirm that Orai1-HaloTag remained fully contained throughout the series and that mobile Orai1-HaloTag channels or autofluorescent aggregates did not enter the ROI. This semi-automated protocol was preferred because fully automated analysis with TrackMate was prone to errors due to the low SNR of dim JF646-BAPTA signals.

All JF646-BAPTA signals were corrected for background fluorescence. Because background varied across the cell (∼1-3 c.u. for 1 mW laser power and 33 ms exposure), we used local values for correction rather than a single cell-wide average. Local backgrounds were measured next to each Orai1-HaloTag ROI from the average intensity projection of the time series and subtracted from the value of the Orai1-HaloTag ROI.

### Identification of gating events

Background-corrected time series data from ImageJ were analyzed using custom routines written in Igor Pro 8. To distinguish true channel gating events from dye blinking, we applied several criteria for selecting traces for analysis:

1) Intensity at +30 mV holding potential is stable and greater than zero.
2) Intensity increases by 3-6 c.u. upon hyperpolarization to -100 mV (under standard conditions of 1 mW power, 33 ms exposure). This encompasses the fluorescence range for a single open channel (Fig. 6C).
3) Minimum SNR of 4.0. This effectively screened out traces with large fluctuations that repeatedly crossed the 50% threshold level, probably due to simultaneous stochastic blinking of multiple dyes.
4) Minimal drift in fluorescence signal. Intensity sometimes slowly fluctuated over seconds. We suspect this is due to movement in the Z direction within the TIRF evanescent field, and such sections of data were excluded from analysis.

The signature criterion for single-channel closures is transitions to the 0-flux level (defined by the fluorescence at +30 mV) rather than background. The likelihood of a blinking event reaching this level declines as the number of dyes per channel increases. Thus, an advantage of the Orai1-HaloTag system is that it is hexameric and can be efficiently labeled with JF646-BAPTA. To identify gating transitions, the closed-state intensity was defined as the mean fluorescence at +30 mV, whereas the open-state intensity was estimated by eye as the maximal average fluorescence at −100 mV, typically ∼2-fold higher than the closed level. Transitions between open and closed states were detected using a 50% threshold between these levels, yielding an idealized binary trace from which open and closed dwell times were extracted. Open probability (P_o_) was calculated as the total fraction of time spent in the open state, based on idealized single-channel traces lasting ≥ 10 s. Dwell-time histograms were fit with single- and double-exponential functions of the form

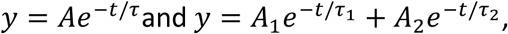

where *y* is the number of events per bin, *A* terms are amplitudes, and *τ* values are time constants. Association and dissociation kinetics were fit using

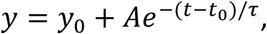

where *y* is signal intensity, *y*_0_ is the baseline offset, *A* is the amplitude, *t* is time with offset *t_0_*, and *τ* is the characteristic time constant.

## ACKNOWLEDGEMENTS

We thank members of the Lewis lab for helpful discussions, Mohamed Trebak (Univ. of Pittsburgh) for generously supplying the Orai1/2/3 triple-knockout HEK293 cell line, Luke Lavis (Janelia Research Campus, HHMI) for the kind gift of JF646-BAPTA/AM, and Miriam Goodman (Stanford) for the use of a micromanipulator mount for the Patch-TIRF experiments. This work was supported by NIH training grant T32MH20016 (S.D.) and by NIH grant R37GM45374 and R35GM149305, a Stanford Discovery Innovation award, and the Mathers Charitable Foundation (R.S.L.).

---

**Figure S1.**
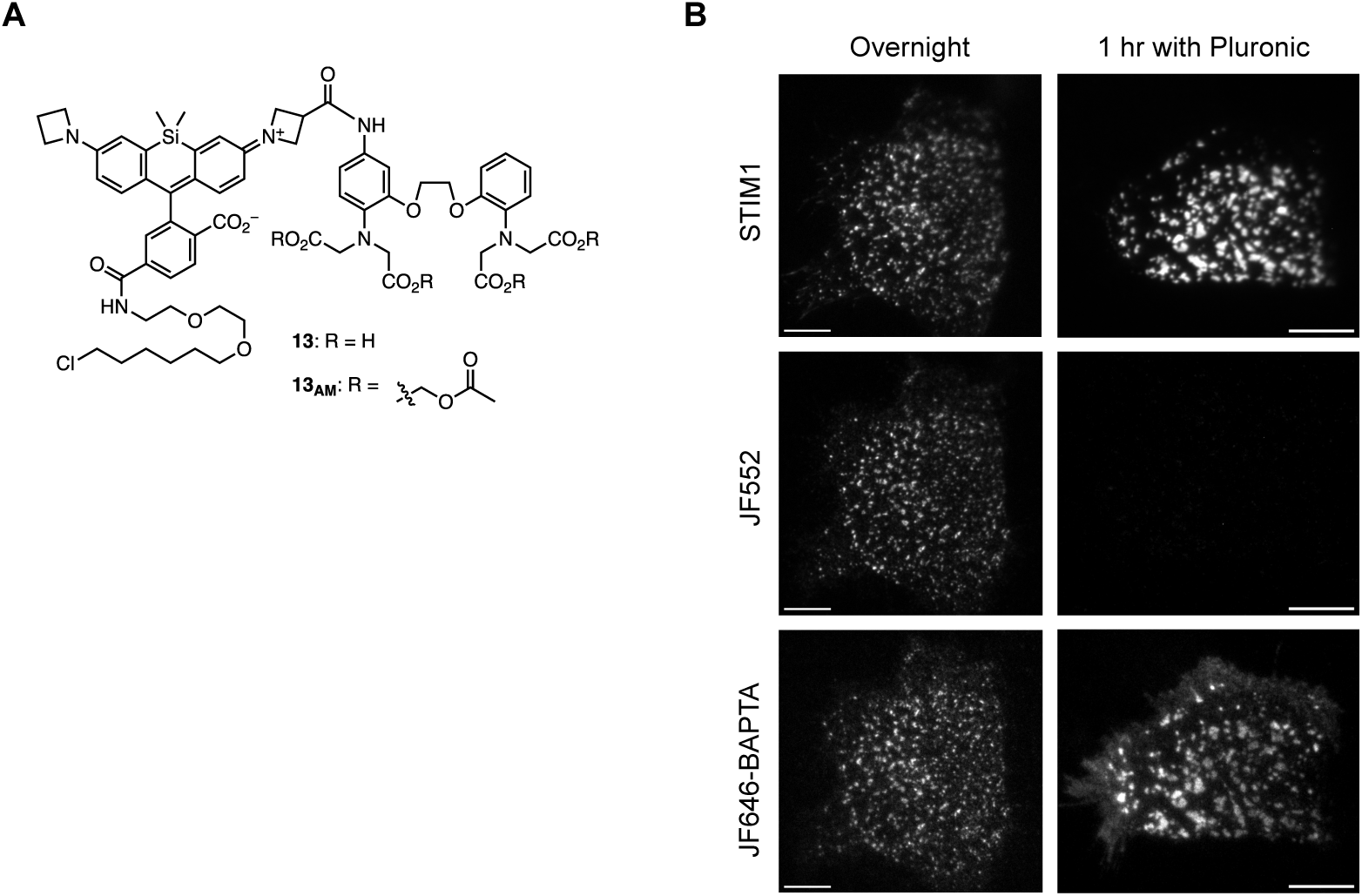
A method for labeling Orai1-HaloTag with the Ca^2+^-sensitive fluor JF646-BAPTA. **(A)** Structure of JF646-BAPTA showing the AM esterified and de-esterified forms (Deo et al., 2019). **(B)** Testing for JF646-BAPTA labeling efficiency. HEK 293 cells were transfected with mCh-STIM1 and Orai1-HaloTag and treated with 1 µM JF646-BAPTA overnight (*left*) or for 1 h in the presence of 0.02% pluronic F-127 (*right column*), followed by exposure to 100 nM JF-552 for 10 min to label unoccupied HaloTag sites. TIRF images of the cell footprints after TG treatment show that pluronic F-127 is needed to achieve complete labeling with JF646-BAPTA. Bar=10 µm.

**Figure S2.**
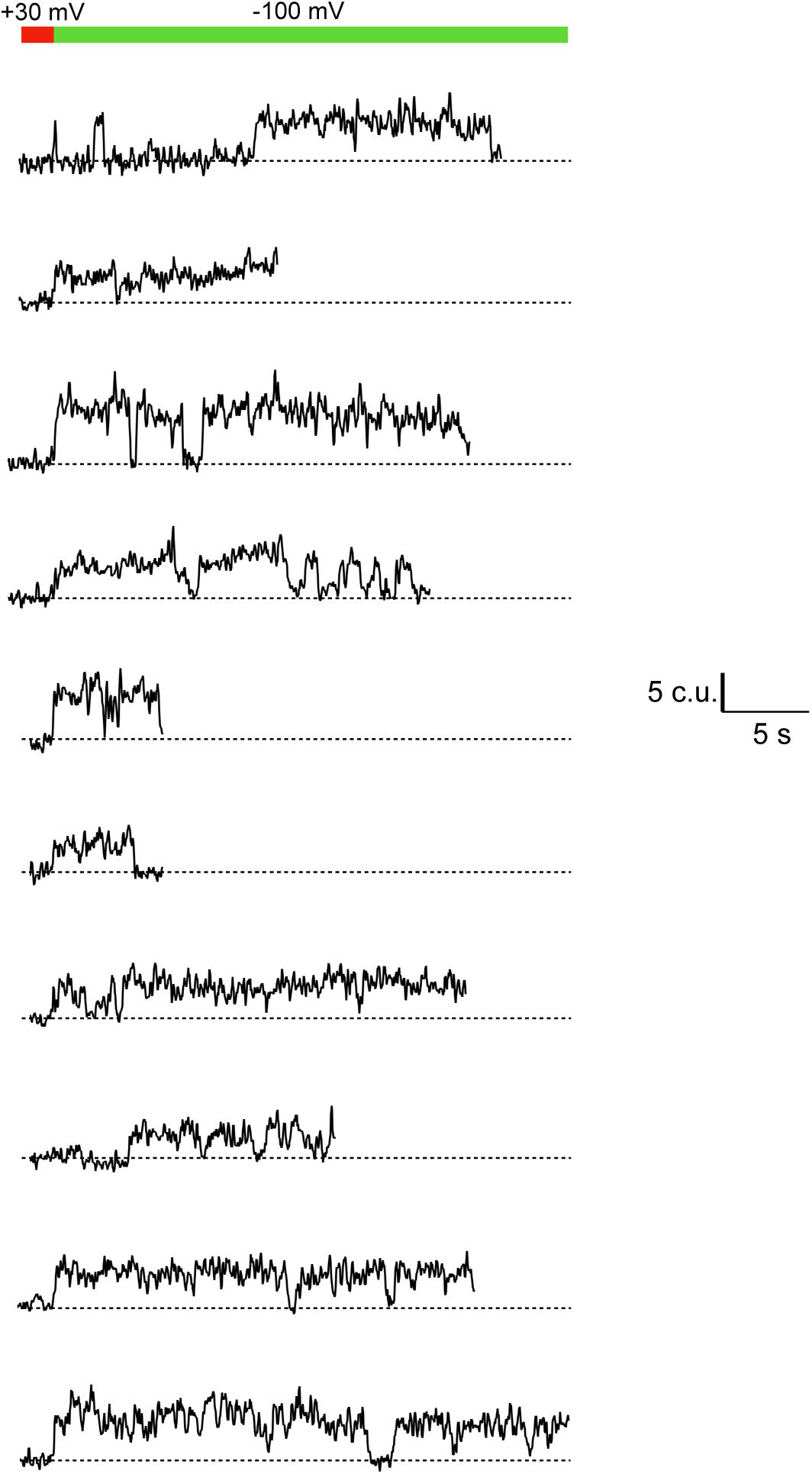
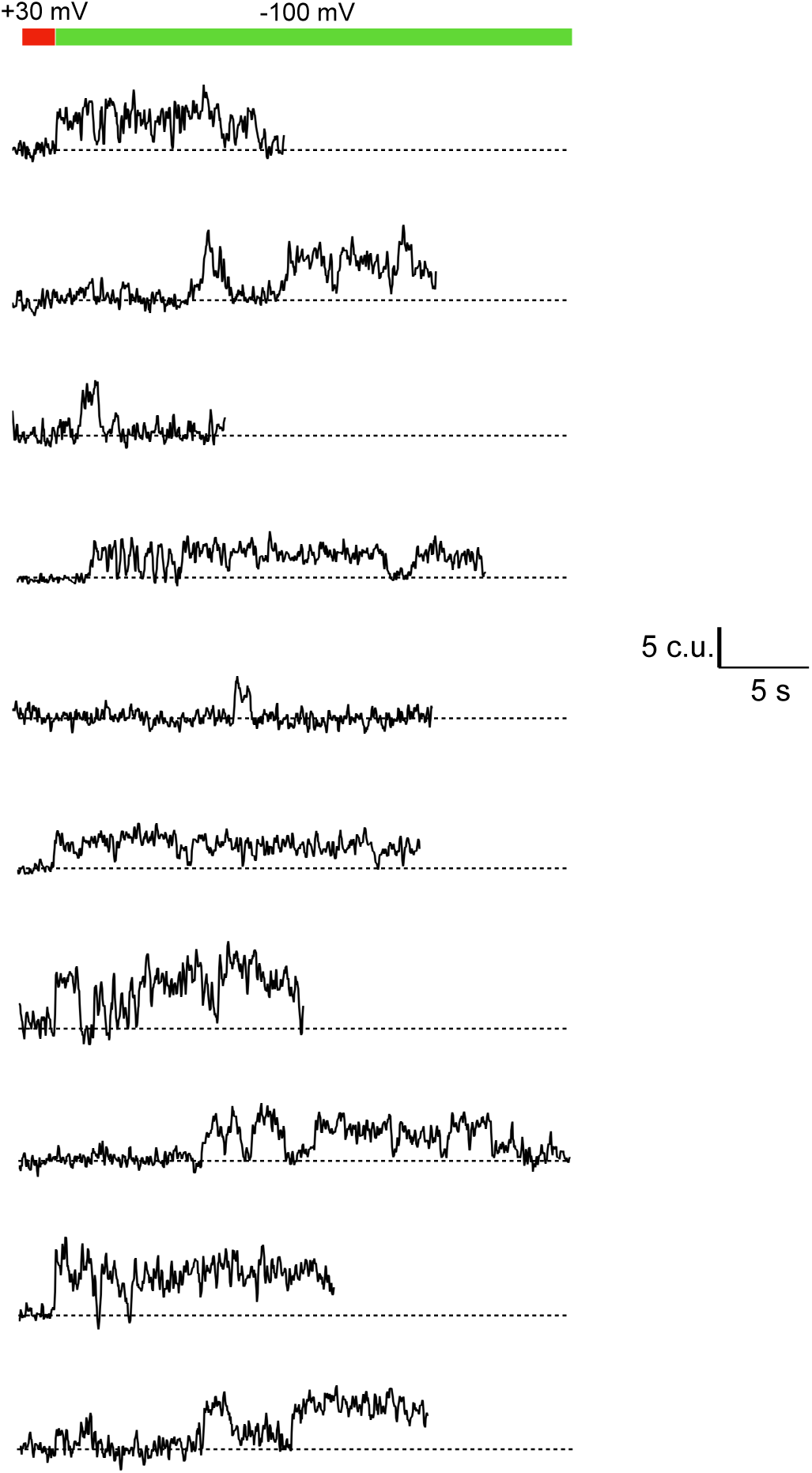
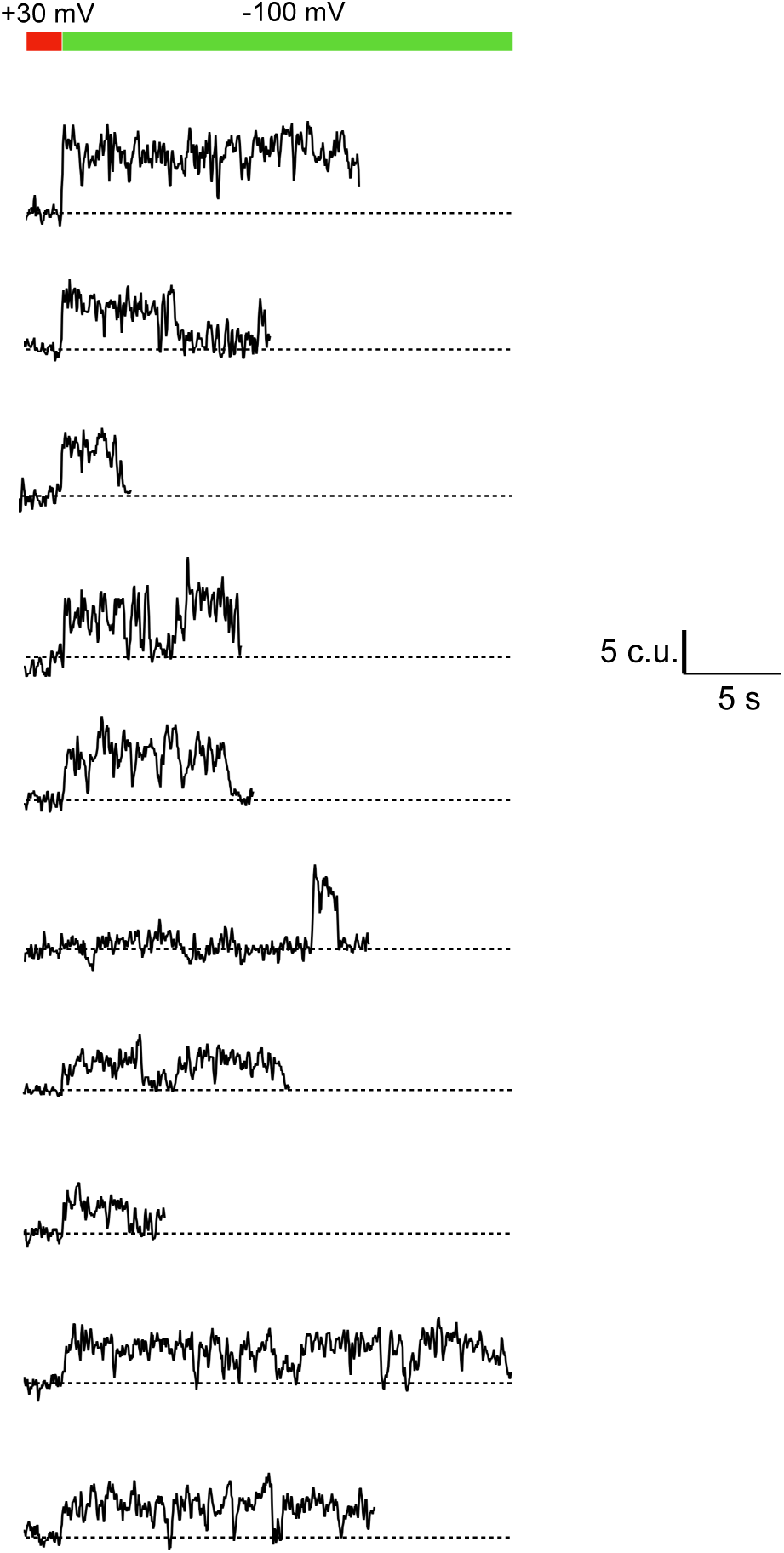
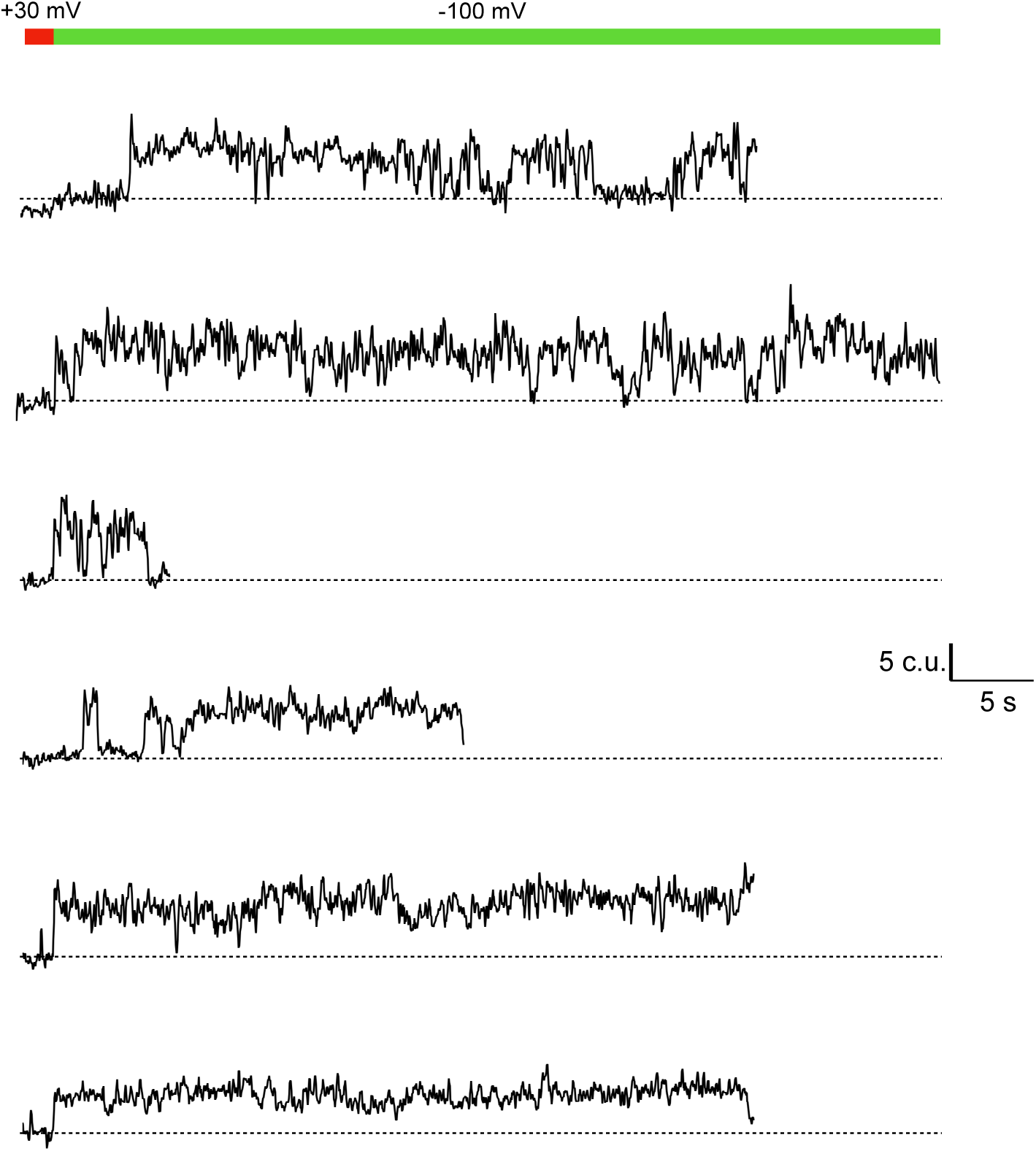
Single-channel optical recordings analyzed for the dwell time histograms. 36 recordings were selected for analysis based on the criteria outlined in Methods. Dashed lines indicate the intensity at +30 mV, as an indicator of the expected closed channel intensity. 33 ms sampling, 1 mW laser power, 3 frame boxcar averaging.

